# OST Catalytic Subunit Redundancy Enables Therapeutic Targeting of N-Glycosylation

**DOI:** 10.1101/2024.12.03.626593

**Authors:** Marta Baro, Hojin Lee, Vanessa Kelley, Rongliang Lou, Chatchai Phoomak, Katerina Politi, Caroline J. Zeiss, Michael Van Zandt, Joseph N. Contessa

## Abstract

Protein asparagine (N)-glycosylation, which promotes folding and trafficking of cell surface receptors such as the EGFR, has not been considered a viable target in oncology due to the essential and non-redundant enzymatic activities required for glycan synthesis and transfer. In mammals an exception to this rule is the presence of the oligosaccharyltransferase (OST) catalytic subunit paralogs, STT3A and STT3B. Here we delineate the chemical biology of OST inhibitors and develop an approach for limited inhibition of N-glycosylation optimized for downstream effects on EGFR. Small molecules with enhanced pharmacokinetic properties and preferences for STT3A or STT3B were synthesized, characterized *in vitro*, and advanced to *in vivo* testing. The lead from this series, NGI-189, causes tumor regression or growth delay of patient derived and TKI resistant EGFR-mutant lung cancer xenografts without toxicity. Together these results suggest that bioavailable OST inhibitors can be developed as therapeutic agents for oncology.

**SUMMARY:** A mechanistic approach for partial inhibition of N-glycosylation delivers small molecules with anti-tumor activity in EGFR mutant NSCLC.

## INTRODUCTION

Post-translational modifications (PTMs) regulate proteins that drive tumor cell proliferation and survival and are frequently targeted in oncology. Successful therapeutic paradigms that alter PTMs include pharmacologic inhibitors of both kinases and histone deacetylases (*1, 2*). These approaches either reduce serine, threonine, and tyrosine phosphorylation to regulate signal transduction programs, or increase histone lysine acetylation to affect gene expression and chromatin structure. Ultimately, however, translation of these strategies to the clinic is accomplished by the design of small molecules that target individual or discrete protein families and thus affect a limited subset of dependent PTMs. Partial pharmacologic inhibition of PTMs thus enables regulation of proteins vital for tumor growth but without the broad effects that cause indiscriminate cellular toxicity.

Asparagine (N)-glycosylation is an endoplasmic reticulum (ER) PTM that promotes proper protein folding through transfer of a 14 carbohydrate-linked core oligosaccharide to an NXT/S/C sequence (where X can be any amino acid except proline), or sequon. This PTM promotes proper protein folding by guiding interaction with the ER’s calreticulin-calnexin quality control cycle (*3*). N-glycosylated proteins that attain mature conformations are then exported to the Golgi apparatus where the glycan undergoes further modifications that guide sorting to the plasma membrane or other organelles within the cell (*4*). Pharmacologic inhibition of N-glycosylation, however, has not been achievable in therapeutic settings due to the toxicity associated with generalized inhibition of this PTM. Nonetheless, a strategy for partial inhibition of N-glycosylation would be valuable, as cell surface glycoproteins that regulate tumor programs such as cell growth (*eg* EGFR (*5, 6*)), angiogenesis (*eg* VEGFR (*7, 8*)), and immune surveillance (*eg* PD-1 (*9, 10*)) have been identified and validated as targets in oncology.

Enhanced EGFR signaling is a frequently identified mechanism of oncogenesis in non-small cell lung cancer (NSCLC), and activating kinase domain mutations are common with a range of incidence from 15-50% in western and eastern populations, respectively (*11, 12*). These mutations are clinically actionable, and targeting EGFR dependent tyrosine phosphorylation with kinase inhibitors (TKI) improves overall survival (*13, 14*) and correlates with population level decreases in lung cancer mortality (*15*). EGFR TKI pharmacology has advanced significantly over the past decade. ATP-competitive and reversible first-generation inhibitors, such as erlotinib and gefitinib, have been followed by a second-generation irreversible inhibitor, afatinib, and by third-generation inhibitors like osimertinib, which are both irreversible and preferentially inhibit mutant EGFR (*16*).

The pharmacologic evolution of EGFR TKIs has been a countermeasure to the emergence of therapeutic resistance that results in rapid expansion of cell populations with secondary mutations (*17, 18*). As a whole, resistance mutations enable EGFR ‘bypass’ signaling; either through EGFR kinase domain mutations, enhanced RTK signaling via co-expressed receptors such as ErbB2 and MET (*19, 20*), or through mutations and fusions of downstream oncogenes. In addition, resistance also occurs through mutation-independent mechanisms such as cell phenotype transitions or activation of other pathways that have yet to be characterized at the molecular level (*21*). Thus, although fourth generation TKIs that target additional EGFR resistance mutations are on the horizon, new strategies to block EGFR and RTK network function that do not directly target kinase activity may provide new pathways for improving the outcomes for this patient population.

EGFR is a type I transmembrane receptor that is highly N-glycosylated, with 12 sequons encoded within the amino acid sequence. EGFR N-glycosylation, as well as that of other proteins, is catalyzed by the oligosaccharyltransferase (OST), a multimeric protein complex embedded in the ER membrane. The OST is composed of a single catalytic subunit, either STT3A or STT3B, six regulatory subunits (RPN1, RPN2, OST48, TMEM258, OST4, DAD1), and catalytic subunit-specific auxiliary subunits (DC2, KCP2, and TUSC3 or MAGT1 (*22, 23*). The OST complex that includes STT3A (OST-A) catalyzes co-translational N-glycosylation and those with STT3B (OST-B) preferentially glycosylate sequons adjacent to disulfide bridges, those located in the distal ∼65 amino acids of the c-terminus, and other sites missed by OST-A (*24–26*). OSTs with different catalytic subunits therefore display both unique and partially overlapping enzymatic functions that ensure the fidelity of N-glycosylation for EGFR, co-expressed RTKs, and other glycoproteins.

In this work we develop small molecule inhibitors of OST enzymatic activity in order to demonstrate that N-glycosylation is an attractive and pharmacologically actionable PTM target. We define a mechanism for limited inhibition of N-glycosylation that takes advantage of OST catalytic subunit redundancy and show that, unlike complete loss of glycan precursor biosynthesis, partial inhibition of N-glycosylation is both achievable and tolerable *in vivo*. Using EGFR mutant and EGFR TKI resistant NSCLC models, OST inhibitor therapeutic efficacy is demonstrated and provides the pre-clinical rationale to pharmacologically target this PTM.

## RESULTS

### Synthesis and optimization of small molecule OST inhibitors

To develop inhibition of N-glycosylation as a therapeutic strategy, we initiated a medicinal chemistry campaign to synthesize potent and bioavailable aminobenzosulfonamide small molecule OST inhibitors that would also be suitable for *in vivo* testing. Because the OST is a multi-subunit and membrane embedded complex, isolation of the enzyme for biochemical analysis of inhibitor activity is not practical. Analogs were therefore tested and screened using the cell-based ER-LucT reporter (*27*), a preferred substrate of the STT3A catalytic subunit (*28*), that is activated upon inhibition of N-glycosylation (*29*). The reporter measures the product of enzymatic activity, glycan site occupancy (Fig. 1a), and provides an advantage over isolated enzyme preparations because it requires small molecule membrane permeability and enzymatic inhibition in the oxidizing environment of the ER lumen in order to produce reporter activity and bioluminescence. Dose response quantification of luciferase activity therefore enables calculation of IC_50_ values and reliable comparisons between analogs.

**Figure 1.**
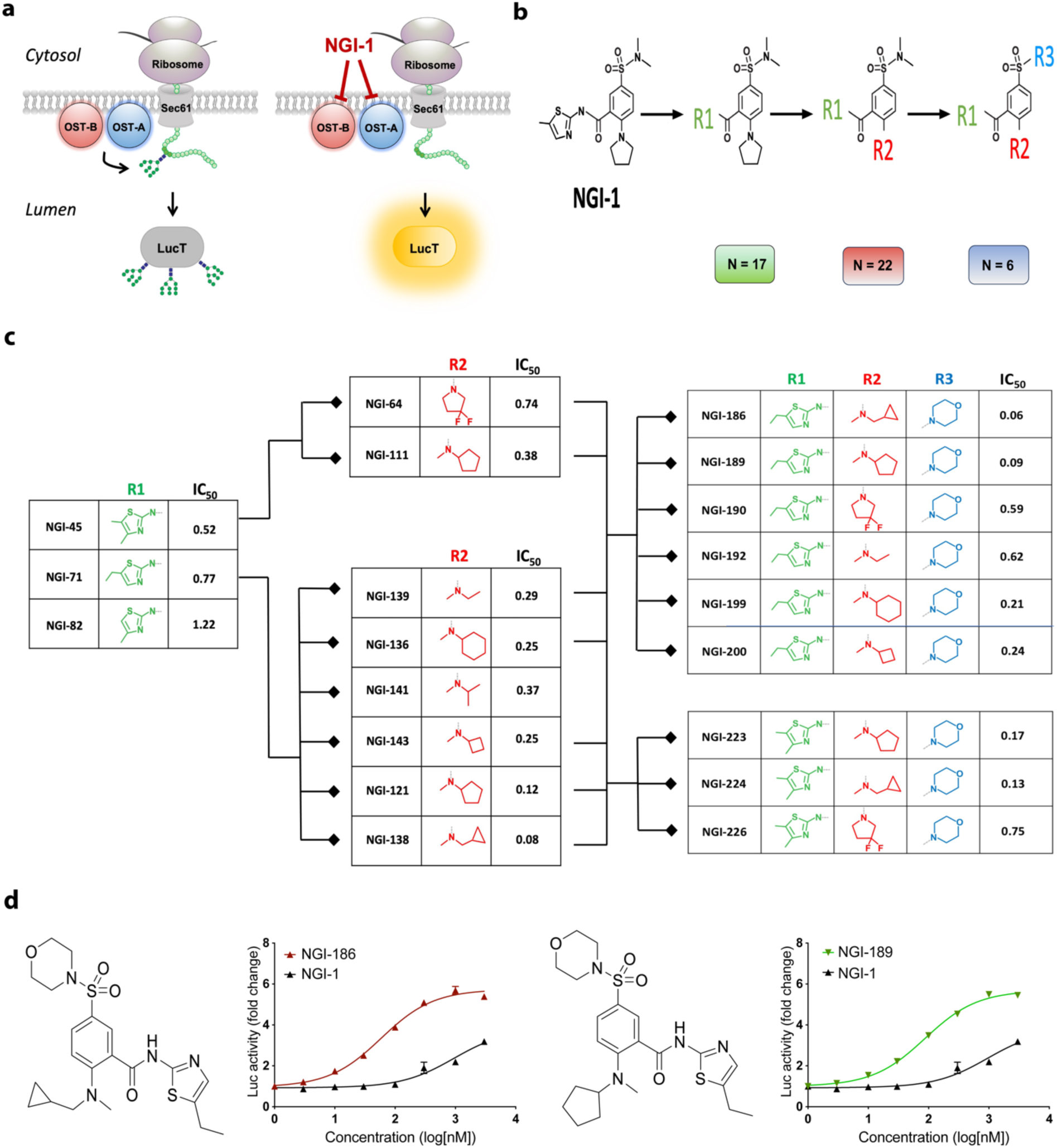
Structure Activity Relationships (SAR) and optimization of OST inhibitor activity. **(a)** Strategy for determination of OST inhibitor in IC_50_ using the ERLucT reporter. Luciferase activity is inactivated under normal conditions but activated upon inhibition of N-glycosylation (*e.g.* NGI-1 treatment). **(b)** Overview for synthesis of OST inhibitors and sequential R1, R2, and R3 group substitutions to the NGI-1 parent molecule. The number of different substitutions for each R group are noted. **(c)** Results of small molecule inhibitor IC_50_ values following serial pharmacophore substitutions. Data represents values from at least two independent experiments performed in triplicate. **(d)** Dose response inhibition of N-glycosylation measured by ER-LucT induced bioluminescence for NGI-1, NGI-186 or NGI-189. Luciferase activity is shown as fold increase for three technical replicates and error bars represent the SEM.

The overall strategy for OST inhibitor optimization through serial R group substitutions is presented in Figure 1b. In total, over 200 different analogs were synthesized and 77 displayed OST inhibitory activity. This included single or combinatorial pharmacophore substitutions with generation of unique R1 (n=17), R2 (n=22), or R3 (n=6) pharmacophores. To drive potency, attention was first directed to the methyl-thiazole group (R1), and it was determined that either 5-ethyl thiazole or 4,5 methyl thiazole substitutions enhance activity (Fig. 1c; green). These improvements were then advanced for further optimizations centered on the pyrrolidine moiety (R2; red). Here, addition of a difluoro group to the 2-position or substitution with various methyl tertiary amines showed further activity enhancement. Of note, methyl-cyclopropyl and cyclopentyl group substitutions enhanced potency by ∼10 fold, with IC_50_ values of 80 and 120nM, respectively. Finally, the sulfonamide group (R3; blue) was substituted by a morpholine-sulfonomide to make additional contributions to potency and anticipated improvements for solubility. A summary of structure activity relationships for twenty analogs with IC_50_ values equal to or better than the NGI-1 lead compound are reported (Fig. 1c), along with the complete dose response activity profiles of the most potent analogs NGI-186 and - 189 (Fig. 1d).

### Defining N-glycosylation dependent and independent EGFR addiction

To identify OST inhibitors that reduce NSCLC proliferation by disrupting EGFR function, we designed a cell based counterscreen with N-glycosylation independent EGFR signaling. This strategy preserves EGFR signaling despite loss of N-glycosylation and is predicted to rescue the EGFR dependent effects of OST inhibitors and to validate the cellular mechanism for reduced proliferation. N-glycosylation independent EGFR signaling was achieved through expression if an EGFR chimera with the extracellular and transmembrane domains of CD8, chosen because it does not require N-glycosylation and spontaneously dimerizes, to the intracellular domain of the EGFR engineered to harbor the L858R kinase domain activation mutation and the C797S osimertinib resistance mutation (Fig. 2a). This chimera, termed CD8-EGFR-CL, is predicted to be rescue cell from OST inhibitors that block EGFR signaling and therefore guide selection of small molecules with the desired biological effect.

**Figure 2.**
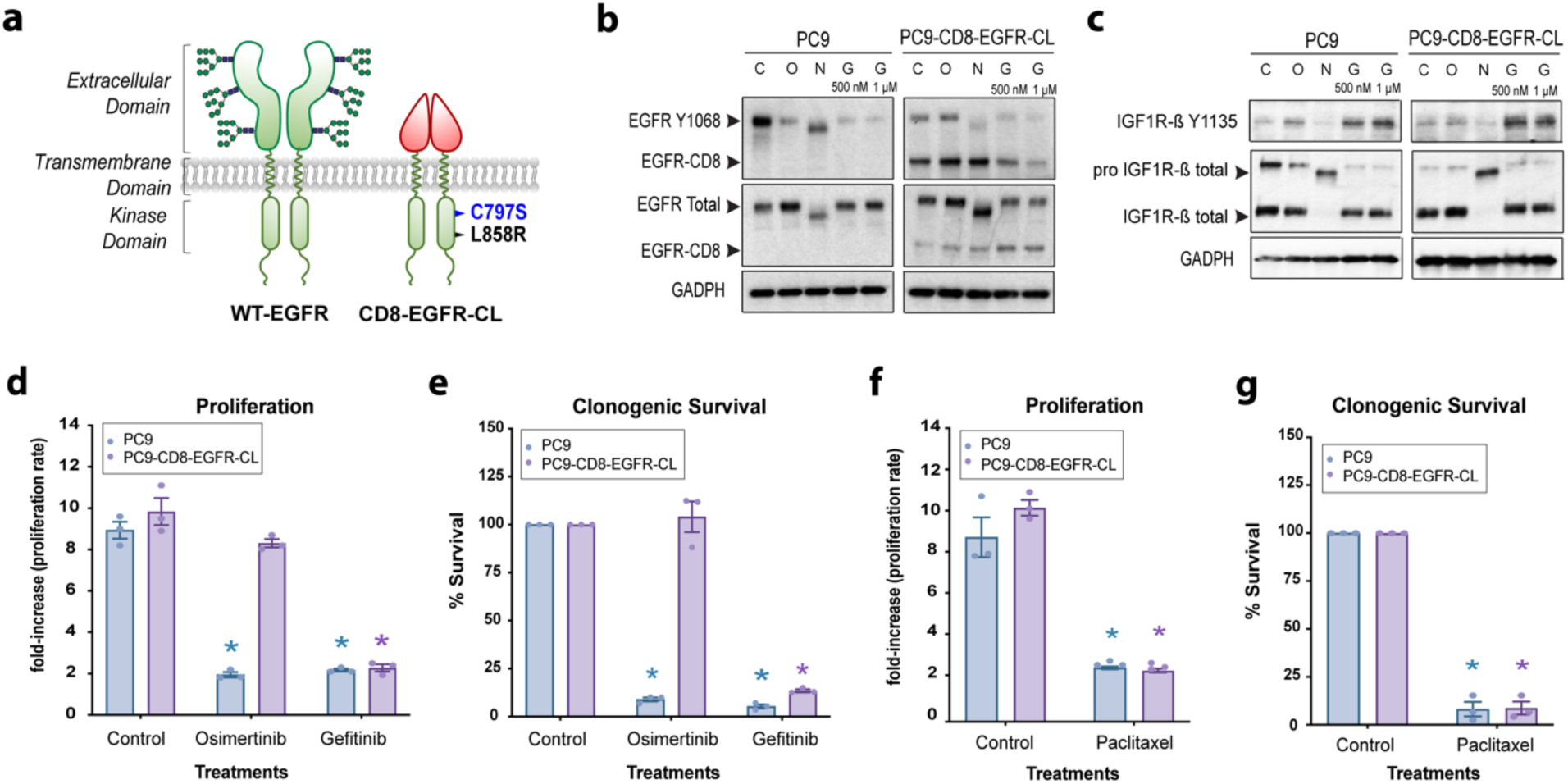
N-glycosylation independent EGFR addiction. **(a)** Cartoon depicting the CD8-EGFR-CL chimeric receptor, which undergoes spontaneous dimerization and harbors the activating L858R mutation and C797S osimertinib resistance mutation. **(b-c)** Western blots of total EGFR and phospho Y1068, or IGF1R-ß and phospho Y1135 in parental and CD8-EGFR-CL expressing PC9 cell lines treated with vehicle control (C), 1 μM Osimertinib (O), 10 μM of NGI-1 (N), or 500 nM or 1 μM of Gefitinib (G) for 48 hours. GAPDH was used as a loading control. Arrows show the EGFR and CD8-EGFR **(b)** or the IGF1R or pro-IGF1R **(c)**. Representative data from two independent experiments are shown. **(d)** Proliferation determined by MTT assay for parental or CD8-EGFR-CL expressing PC9 cells after 5 days of control, 1 μM Osimertinib, or 500 nM Gefitinib treatment. **(e)** Clonogenic survival analysis using the same experimental design as in (d). **(f-g)** The effects of 2.5 nM paclitaxel in MTT (f) or clonogenic survival (g) analysis for PC9 and PC9-CL cells. For all bar graphs results show mean values ± SEM for at least three independent experiments. An * indicates a significant difference (P ≤ 0.05).

In parental EGFR mutant PC9 NSCLC, OST inhibition with NGI-1 reduces phosphorylation of the endogenous EGFR (Fig. 2b). In contrast, CD8-EGFR-CL phosphorylation is not blocked by NGI-1, confirming that the chimera is not dependent on N-glycosylation for activity. CD8-EGFR-CL expression does not upregulate other co-expressed RTKs known to initiate EGFR bypass survival signaling such as IGF-1R (*30, 31*), which remained sensitive to NGI-1 (Fig. 2c). To determine whether CD8-EGFR-CL expression sustains EGFR oncogene addiction, the effects of gefitinib and osimertinib were compared in proliferation (Fig. 2d) and colony formation (Fig. 2e) assays. CD8-EGFR-CL expressing cells were resistant to osimertinib (due to the C797S mutation) but sensitive to gefitinib, establishing that oncogenic signaling and EGFR addiction is indeed preserved by the chimera. In contrast no difference in proliferation (Fig. 2f) or colony formation (Fig. 2g) between parental and CD8-EGFR-CL expressing cells was observed for a mechanistically different anti-proliferative agent, paclitaxel, arguing that CD8-EGFR-CL expression does not cause generalized therapeutic resistance. Together these results validate expression of the CD8-EGFR-CL chimera as an effective tool for generating N-glycosylation independent, EGFR signaling and for isolating EGFR dependent effects on tumor cell growth and survival.

### Inhibitor screening in EGFR mutant, N-glycosylation independent NSCLC

OST inhibitors with structural diversity and enhanced IC_50_ values were then selected for testing in the PC9/PC9-CD8-EGFR-CL cell line pair. Inhibitor treatments reduced proliferation from 20-70% in parental cells as measured by MTT assays at 5 days (Fig. 3a). We found that while reductions in proliferation seen in PC9 cells correlated with ER-LucT derived IC_50_ values, the luciferase assay was not entirely predictive. Instead, the effect of each inhibitor on EGFR phosphorylation was a better predictor of anti-proliferative effect (Fig. 3b). For example, NGI-111 (IC_50_ = 380nM) did not reduce EGFR phosphorylation or cause visible molecular weight changes (Fig. 3b). A lack of EGFR molecular weight change was also observed for this inhibitor in HEK293 cells, as well as for the engineered Halo3N glycoprotein that is co-expressed in this cell line (Fig. 3c; (*28*)). Together the results demonstrate the challenges of using a general indicator of N-glycosylation (*ie* ER-LucT) to predict N-glycosylation effects on individual proteins such as the EGFR. These observed differences validate a strategy of developing secondary and tertiary approaches for evaluating OST inhibitors effects in EGFR specific assays as a path towards analog development. For each of the six inhibitors that reduced proliferation by at least 40%, a significant rescue was observed by expression of the CD8-EGFR-CL. In addition, a larger inhibitory effect on PC9 proliferation was significantly correlated with a greater rescue by CD8-EGFR-CL (R=0.92, p < 0.01 (Fig. S1)), consistent with OST inhibitor effects in EGFR mutant NSCLC being primarily mediated by disruption of EGFR glycosylation.

**Figure 3.**
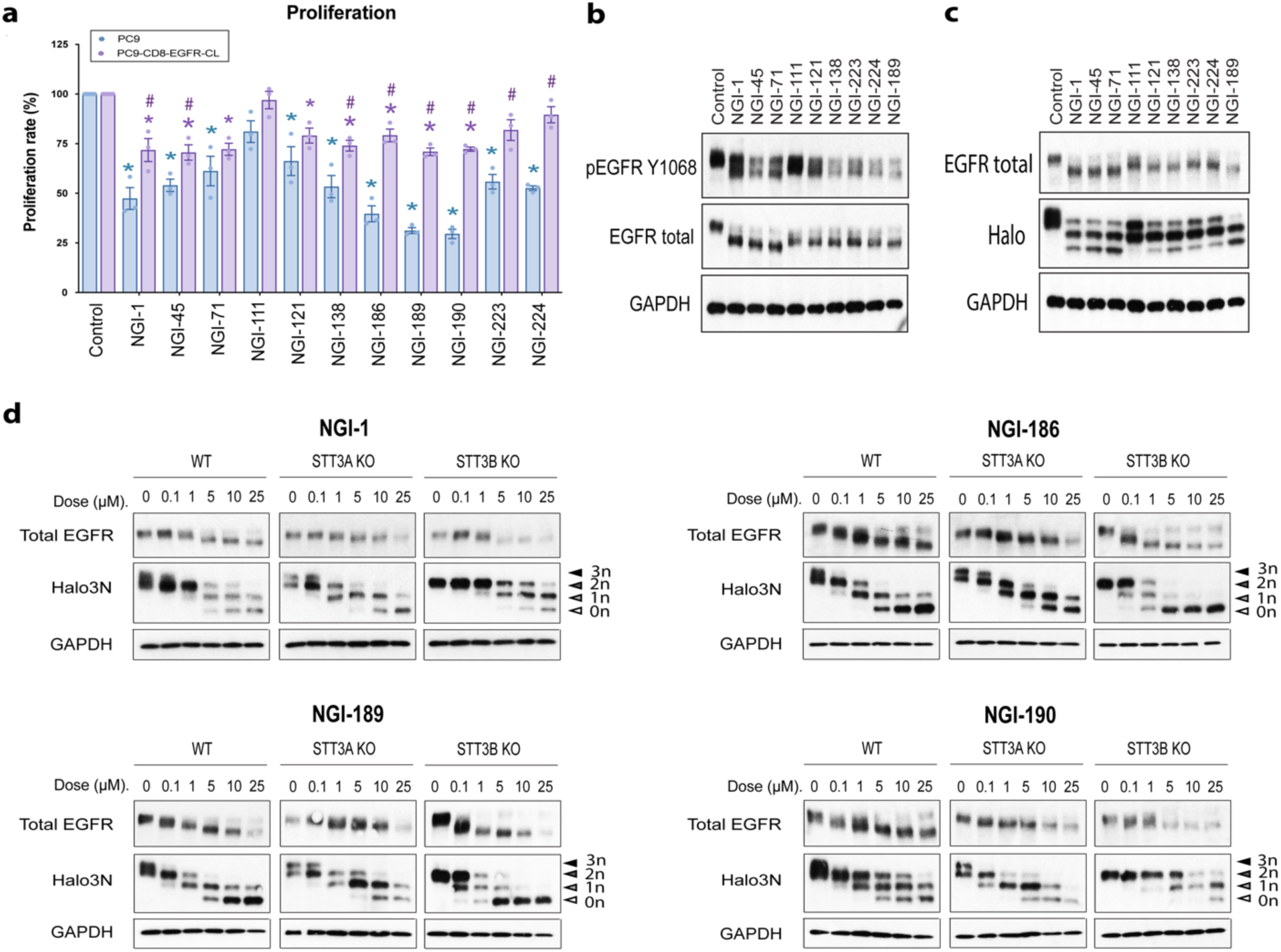
Screen for OST inhibitor activity and catalytic subunit inhibition. **(a)** The effects of OST inhibitors (5 μM) on proliferation in parental and CD8-EGFR-CL expressing PC9 cell lines after 5 days of drug exposure. **(b-c)** Western blots comparing inhibition of EGFR signaling in PC9 **(b)** and reduction of EGFR and Halo3N size in HEK293 parental **(c)** cell lines treated with 5 μM of each inhibitor for 24h. GAPDH was used as a loading control. Representative data from two independent experiments are shown. **(d)** Western blots of HEK293 parental (WT), STT3A KO and STT3B KO cell lines expressing the Halo3N glycoprotein and treated with 0, 0.1, 1, 5, 10 or 25 μM of NGI-1, NGI-186, NGI-189 or NGI-190 for 24 hours. Reduction of EGFR and Halo3N size is caused by reduced N-glycosylation. GAPDH was used as a loading control. Representative data from two independent experiments are shown.

The proliferation results also identified one inhibitor, NGI-190, to have greater effects than what would be predicted by the ER-LucT IC_50_ value (590nM). Because OST-A and OST-B complexes, accomplish high fidelity EGFR N-glycosylation through co- and post-translational transfer of glycans, respectively, we analyzed the effects of NGI-190 and the other two most potent inhibitors (NGI-186, −189) on EGFR and Halo3N glycosylation using HEK293 cells with either STT3A or STT3B knockout. Each inhibitor demonstrated activity against both OST-A and OST-B, and preferential inhibition of OST-A (NGI-186 or NGI-189) or OST-B (NGI-190) was also identified (Fig. 3d). Consistent with IC_50_ analysis calculated by ER-LucT activation (Fig. 2a), NGI-186 and NGI-189 reduce OST-A activity at doses approximately ten times lower than NGI-1 while inhibiting OST-B at similar concentrations to NGI-1. In comparison, NGI-190 is a more potent inhibitor of OST-B but reduces OST-A activity at similar concentrations to NGI-1. These experiments are also notable for the effect of NGI-186 and −189 or NGI-190 at high doses (>10µM). In KO cells complete inhibition of STT3A or STT3B dependent N-glycosylation is achieved. However, in wild type cells with both OST-A and OST-B, complete inhibition cannot be achieved. This finding reveals that the partial inhibition of N-glycosylation caused by this class of inhibitors is due to OST catalytic subunit redundancy. A second important observation is that improved inhibition of either OST-A or OST-B produced greater effects on EGFR N-glycosylation, suggesting that receptor maturation requires sites dependent on both OST-A and OST-B. This result provides a fundamental insight for therapeutic development of OST inhibitors in that biologic activity requires optimization for a target glycoprotein instead of for a specific OST catalytic subunit.

### OST inhibition reduces EGFR Mutant NSCLC survival

The EGFR drives the cell cycle and the effects of the two most potent inhibitors (NGI-186 and −189) on G1 to S progression in PC-9 cells were then evaluated. Indeed these optimized OST inhibitors induced a G1 arrest (Fig. 4a, b), consistent with a more potent inhibition of EGFR phosphorylation (Fig. 4c). The effects of NGI-186 and NGI-189 on EGFR survival signaling were also evaluated, and showed a greater dose-dependent inhibition of AKT, p70 S6K and S6RP phosphorylation (Fig. 4d). In addition, NGI-186 and −189, but not NGI-1, caused induction of the pro-apoptotic protein, Bim. The effects of each OST inhibitor were next tested using colony formation assays to determine changes in survival, and a greater reduction of PC9 clonogenicity with significant rescue by CD8-EGFR-CL expression was observed (Fig. 4e, f). Together these results indicate that optimized SAR translates into greater impairment of EGFR dependent signaling programs that regulate cell cycle progression, proliferation, and survival.

**Figure 4.**
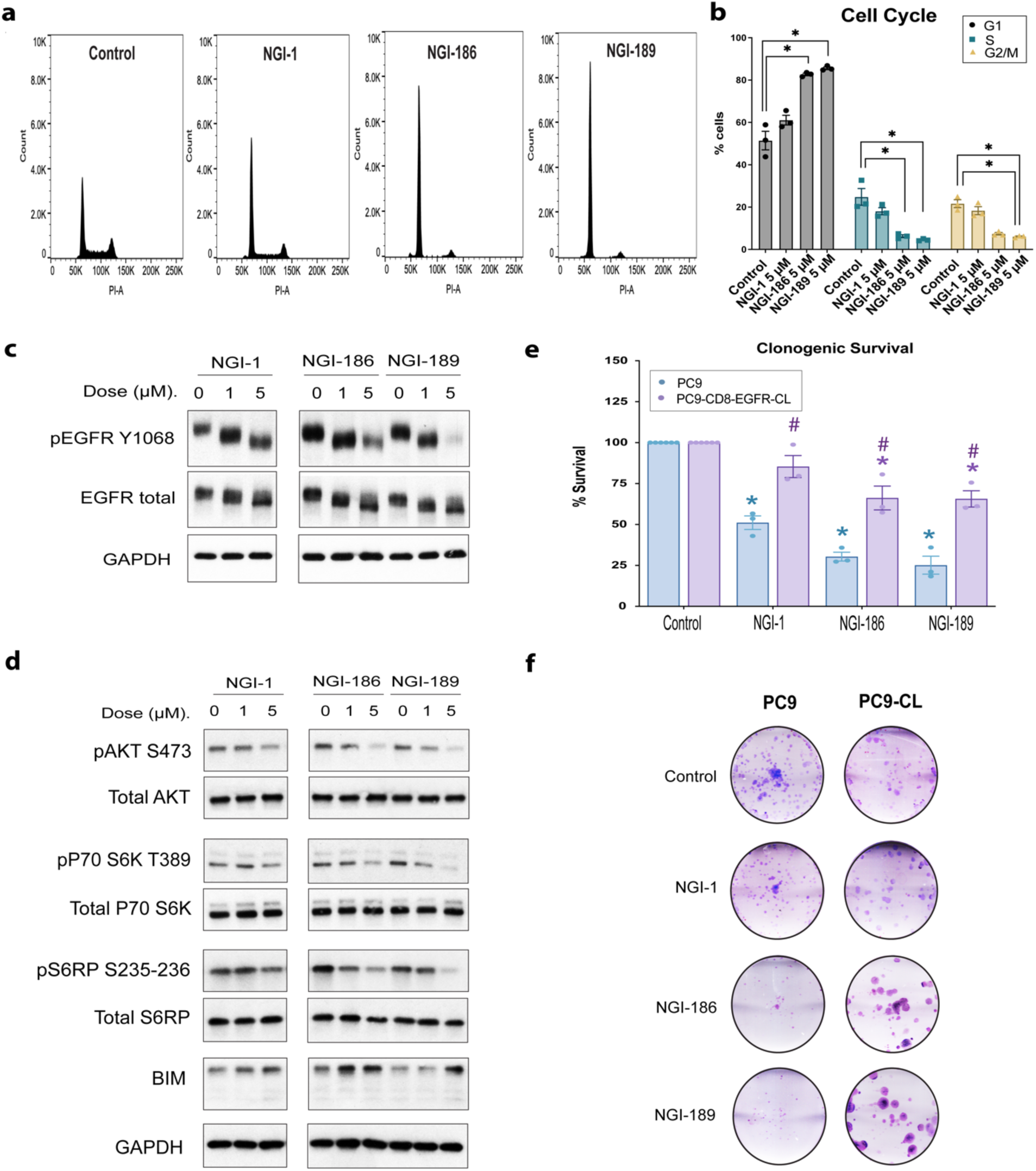
OST inhibitor effects on proliferation and survival in EGFR mutant NSCLC. (a-b) Cell cycle distribution and quantification of PC9 cells treated with vehicle or 5 μM NGI-1, NGI-186 or NGI-189 for 24 hours. **(c-d)** Western blots comparing inhibition of EGFR signaling in PC9 cells line treated with 5 μM of NGI-1, NGI-186, or NGI-189 for 24h. GAPDH was used as a loading control. Representative data from two independent experiments are shown. **(d)** Clonogenic survival of parental and CD8-EGFR-CL expressing PC9 cell lines treated with vehicle, or 5 μM of NGI-1, NGI-186, or NGI-189. **(e)** Representative images of clonogenic survival assays with OST inhibitors. For all bar graphs results show mean values ± SEM for at least three independent experiments. An * indicates a significant difference vs. the control condition (P ≤ 0.05). An # indicates a significant difference between PC9 and PC9-CL value (P ≤ 0.05).

Given the demonstrated efficacy and on-target mechanism of signaling inhibition in PC9 cells, the effects of NGI-186 and −189 were next tested in a panel of additional EGFR mutant NSCLC cell lines: H3255 (EGFR L858R), HCC-4006 (EGFR del747-749, A750P), or HCC-2935 (EGFR del746-751, S752I). In each cell type, optimized analogs caused greater inhibition of signaling through EGFR, Akt, p70 S6K, and S6RP (Fig. 5a, b), with enhanced induction of Bim. In these NSCLC cell lines, OST inhibitors induced a more modest increase in G1 cell cycle arrest, but treatment also caused the appearance of a sub-G1 population of cells, consistent with induction of cell death (Fig. 5c). This effect was also accompanied by enhanced cell killing as measured by clonogenic survival analysis (Fig. 5d). Together these results show that OST inhibitor pharmacology optimized to target the EGFR achieves *in vitro* anti-tumor effects in EGFR mutant NSCLC.

**Figure 5.**
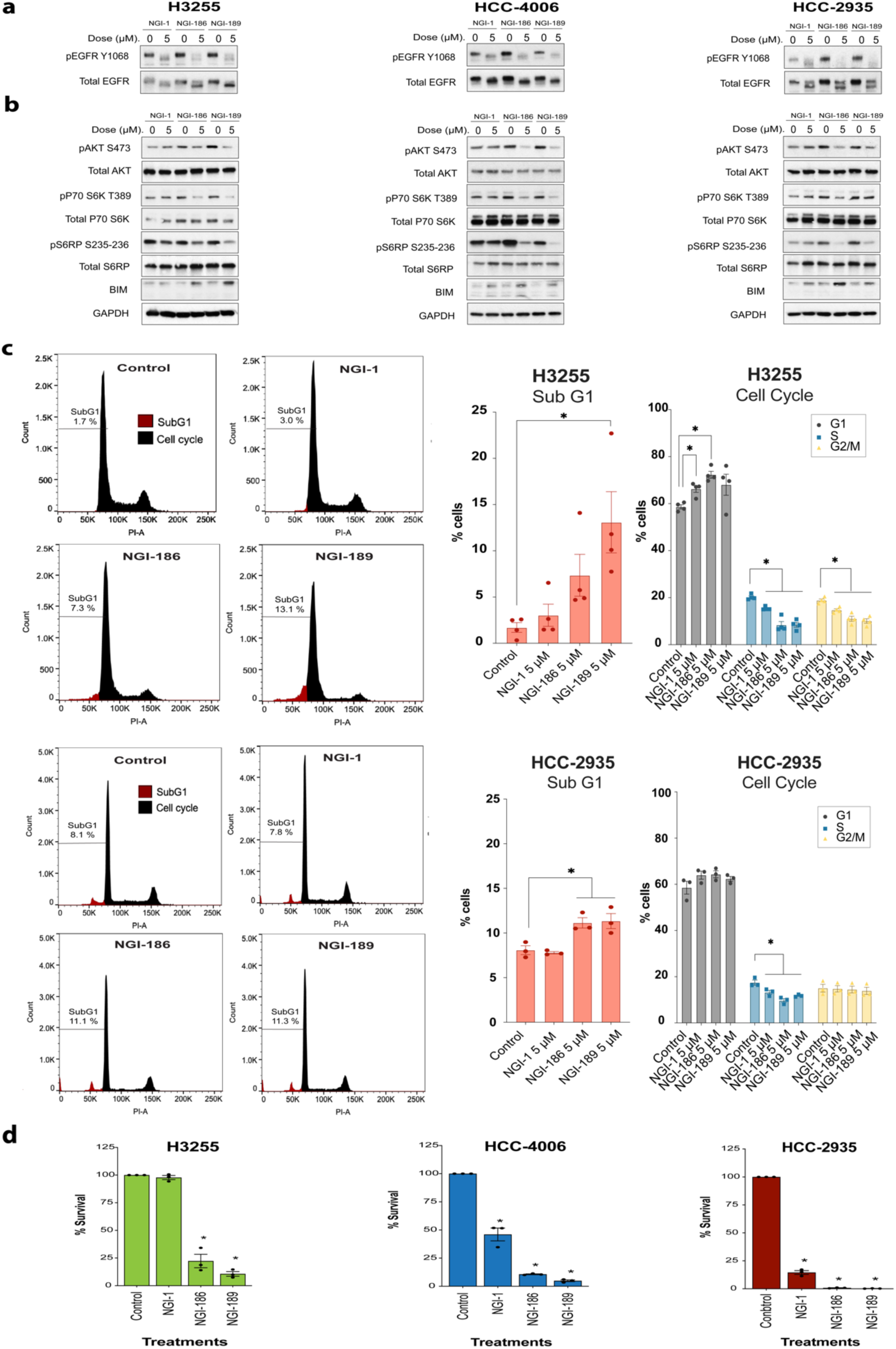
OST inhibition blocks EGFR dependent signaling in EGFR mutant NSCLC. (a-b) Western blots comparing inhibition of EGFR signaling in H3255, HCC-4006 and HCC-2935 NSCLC cell lines treated with 5 μM of NGI-1, NGI-186, or NGI-189 for 24h. GAPDH was used as a loading control. Representative data from two independent experiments are shown. **(c)** Cell cycle distribution and quantification for H3255 or HCC-2935 cells treated with vehicle or 5 μM of NGI-1, NGI-186 or NGI-189 for 24 hours are shown. **(d)** Clonogenic survival of H3255, HCC-4006 and HCC-2935 NSCLC cell lines treated with vehicle or 5 μM of NGI-1, NGI-186 or NGI-189. Bar graph results show the mean values ± SE for three independent experiments for each cell line. An * indicates a significant difference (P ≤ 0.05).

### Tolerability of OST inhibition *in vivo*

Over the past 45 years, inhibition of N-glycosylation *in vivo* has only been achievable through disruption of glycan precursor synthesis with tunicamycin. This natural product blocks all N-glycosylation and is highly toxic. However, with the redundant activity of OST catalytic subunit paralogs, we hypothesized that aminobezamidosulfonamide OST inhibitors with a therapeutic window for translation could be identified. Previously NGI-1 was shown to have a significant solubility liability that prevented *in vivo* testing (*10, 32*). However, with advances in medicinal chemistry, we proceeded to evaluate the possibility of systemic drug administration and *in vivo* testing. As a first step we measured induction of bioluminescence at 0, 6 and 24h following intraperitoneal (*i.p.*) vehicle or NGI-186 (10mg/kg) treatment in mice bearing MDA-MB-231-ERLucT xenografts. NGI-186 significantly induced bioluminescence at 24 hours (p=0.035; Fig. 6a, b), indicating inhibition of N-glycosylation within the tumor and the feasibility of systemic administration. To select an inhibitor for advancement, the solubility of NGI-186 and NGI-189 was then determined as described in Materials and Methods with values of 14.2 and 23.8 µM, respectively. Given the enhanced solubility and otherwise similar characteristics, NGI-189 was advanced for all further preclinical tumor studies.

**Figure 6.**
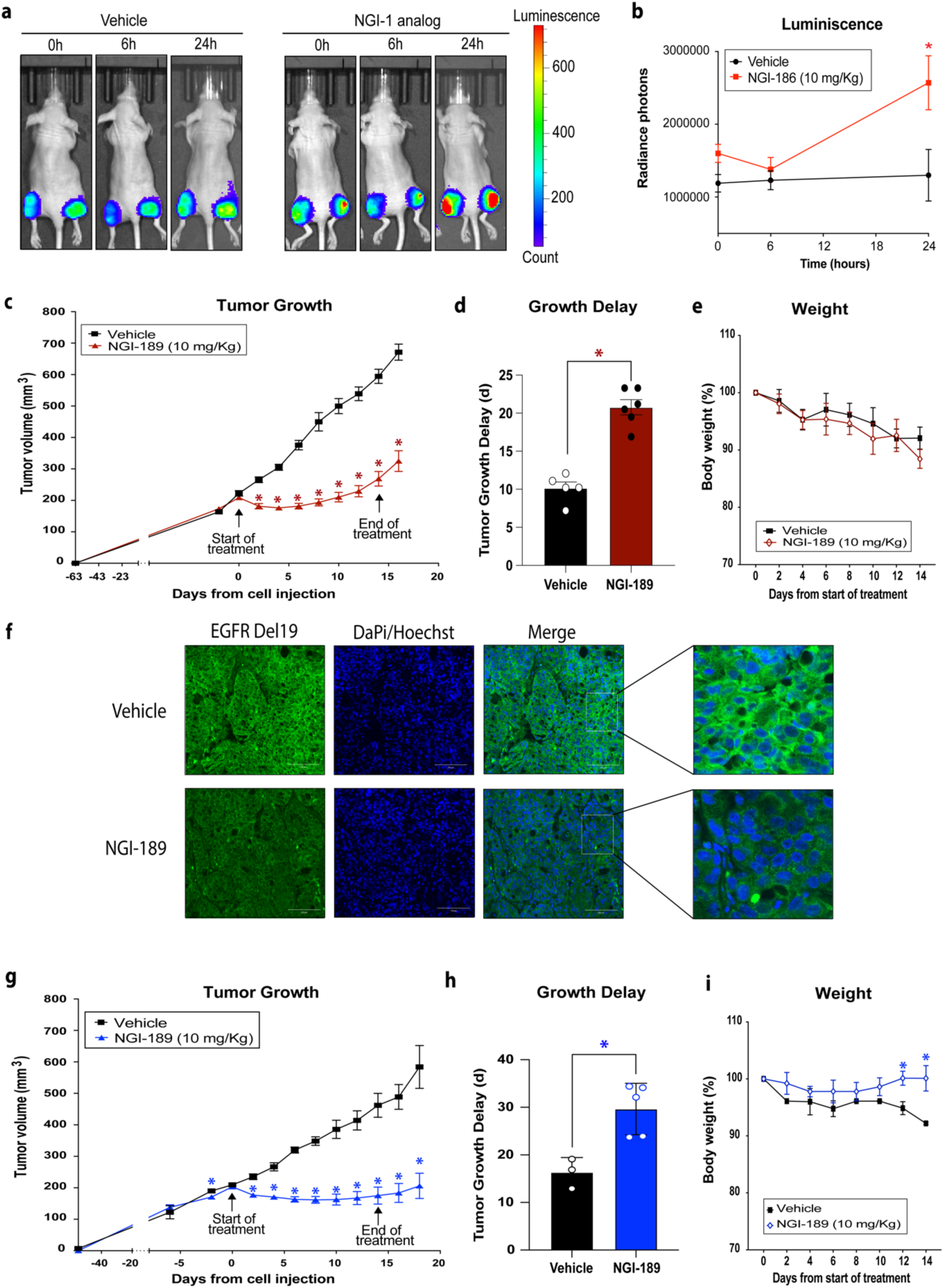
OST inhibitor bioavailability and efficacy in EGFR mutant PDXs. **(a)** *In vivo* imaging of MDA-MB-231 ER-LucT xenografts treated with vehicle or NGI-186 (10 mg/kg). Representative images at 6 and 24 hours demonstrate induction of bioluminescence. **(b)** Average values for bioluminescence with standard error for vehicle (n = 4) or NGI-186 (n = 8) over 24 hours. **(c)** YU-006 average PDX tumor growth ± SE following treatment with vehicle (n = 5) or 10 mg/kg of NGI-189 (n = 6). Treatment was delivered by *i.p.* injection every other day for 8 doses. **(d)** Quantification of tumor growth delay with NGI-189 treatment (days to reach 500 mm^3^). **(e)** Average weight of vehicle or NGI-189 groups during for mice bearing YU-006 PDXs. **(f)** Representative images of EGFR immuno-fluorescence in YU-006 xenografts treated with 3 doses of vehicle (n = 2) or 10 mg/kg of NGI-189 (n = 2). **(g)** YU-010 average PDX tumor growth ± SE following treatment with vehicle (n = 3) or 10 mg/kg of NGI-189 (n = 5). **(h)** Quantification of tumor growth delay and **(i)** Average mouse weight for YU-010 bearing mice as described above. An * indicates a significant difference of vehicle vs NGI-189 (P ≤ 0.05).

Inhibition of all N-glycosylation is toxic in vivo, but the effects of partial disruption of this process are unknown. We therefore performed blood counts, chemistries, and cytopathology for mice treated *i.p.* with NGI-189 or vehicle every other day for a total of three doses with blood and organ sample collection 24 hours following the last dose. The results show no significant differences between vehicle and NGI-189 treatment groups for organ cytopathology (Table S1), for blood counts (Table S2), or for chemistries (Table S2) and overall only minor abnormalities consistent with age related changes were observed. These results provide the first evidence that systemic delivery of an OST inhibitor is tolerable *in vivo* and demonstrate a strong rationale for advancing NGI-189 to efficacy testing in rodent tumor models.

### OST inhibition reduces growth of EGFR mutant xenografts

To assess therapeutic efficacy, NGI-189 was tested in EGFR mutant NSCLC patient derived xenograft (PDX) models, YU-006 and YU-010, which harbor EGFR kinase domain activating deletions in exon 19 (*33*). Established and palpable PDX tumors that grew for up to 9 weeks were randomized to NGI-189 (10mg/kg) or vehicle treatment groups after reaching a size of 200 mm^3^. In YU-006, NGI-189 treatment caused a significant tumor growth delay compared to vehicle-treated tumors (Fig. 6c). By Day 8, vehicle treated YU-006 xenografts more than doubled in size to 450 ± 29 mm^3^, whereas NGI-189 treated tumors showed no significant progression from the beginning of treatment with an average volume of 193 ± 11 mm^3^ (p < 0.001). The time to reach 500 mm^3^ was approximately doubled at 10.2 vs 20.8 days, respectively (Fig. 6d; p< 0.0001). Treatment with NGI-189 was well tolerated and small variations in weight were similar to vehicle treated mice (Fig. 6e). EGFR immunofluorescence for EGFR in YU-006 PDX bearing mice with larger tumors (∼600-700 mm^3^; that also regressed Fig. S2) treated every other day with NGI-189 or vehicle control was also performed. These tumors showed a reduction of EGFR protein levels consistent with decreased receptor maturation following OST inhibition (Fig. 6f).

Significant growth delay was also observed for YU-010 PDXs. At day 12 after the start of treatment, average tumor volumes for vehicle treated mice more than doubled to 414 ± 30 mm^3^, but xenografts did not progress in the NGI-189 treated group, with a significantly smaller average volume of 167 ± 21 mm^3^ (Fig. 6g; p= 0.003). The time to reach 500 mm^3^ was also approximately doubled at 16.3 vs 29.6 days, respectively (Fig. 6h; p= 0.005). OST inhibition was again well tolerated with NGI-189 treated mice having significantly higher weight at the end of treatment, which we interpret to reflect a reduction of tumor burden (Fig. 6i).

EGFR mutant NSCLC eventually develops TKI therapeutic resistance, often through bypass RTK signaling, and presents a significant therapeutic challenge. Because RTK glycoproteins are among the most highly N-glycosylated proteins (*34*), and are also sensitive to OST inhibition (*35*), we hypothesized that these resistance mechanisms would also be susceptible to OST inhibition *in vivo*. Effects of OST inhibition on osimertinib resistant (OR) H1975 cells, which harbor EGFR L858R and T790M mutations and have been reported to upregulate PTK7 signaling as a resistance mechanism (*36*), were therefore examined. Western blots confirmed that H1975-OR cells indeed demonstrate enhanced expression of PTK7, and show that OST inhibition reduces N-glycosylation of PTK7 and represses PTK7/STAT3 signaling (Fig. 7a). Interestingly, we also observed PTK7 expression in both the parental HCC-827 cells, which harbor an EGFR deletion 19 mutant (E746-A750), and the MET amplified and gefitinib resistant (GR) HCC-827 subline (*37*). OST inhibition with NGI-1, NGI-186, or −189 reduced both PTK7 and MET N-glycosylation as well as downstream STAT3 signaling (Fig. 7b) in this cell type. Consistent with reduced IGF-1R signaling *in vitro* (Fig. 3c), these results provide additional evidence that OST inhibition disrupts both EGFR and its bypass signaling mechanisms in NSCLC.

**Figure 7.**
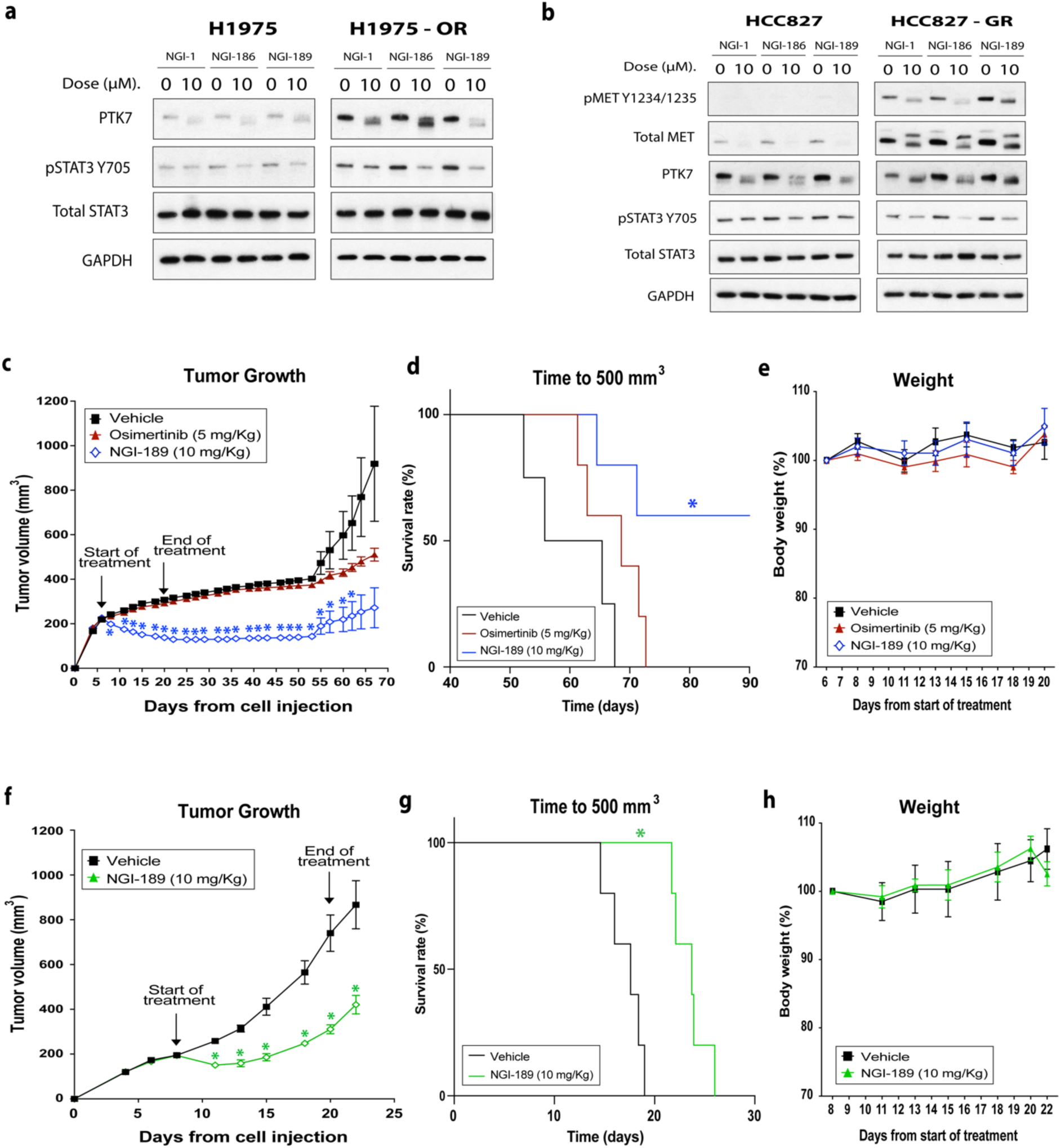
OST inhibitor efficacy in EGFR mutant and TKI resistant xenografts. (a-. **b)** Western blots of H1975, H1975-OR, HCC827 and HCC827-GR NSCLC cell lines treated with vehicle or 10 μM of NGI-1, NGI-186 or NGI-189 for 24 hours. Representative data from two independent experiments are shown. **(c)** Mean H1975-OR xenograft tumor growth following treatment with vehicle (n = 5), osimertinib *p.o.* 5 mg/kg daily (n = 5), or NGI-189 *i.p.* 10 mg/kg (n = 5) every other day. **(d)** Quantification of survival rate (time to reach 500 mm^3^) for vehicle (n = 4), Osimertinib (n = 5), or NGI-189 (n = 5) treatment in H1975-OR tumors. **(e)** Average weight of vehicle or NGI-189 treated mice with H1975-OR xenografts are shown. **(f)** HCC827-GR xenograft tumor growth following treatment with vehicle (n = 5) or 10 mg/kg of NGI-189 (n = 5) delivered *i.p.* every other day for 8 doses. **(g)** Quantification of survival rate (time to reach 500 mm^3^) for vehicle (n = 5) or NGI-189 (n = 5) treatment in HCC827-GR tumors. **(h)** Average weight of vehicle or NGI-189 treated mice from HCC827-GR are shown. Values are the means ± SE and an * indicates a significant difference between vehicle and NGI-189 treatment (p ≤ 0.05).

H1975-OR grow slowly as xenografts due to reduced fitness, and as anticipated, did not respond to osimertinib (5mg/kg daily for 2 weeks). In contrast to osimertinib, NGI-189 caused significant tumor regression (Fig. 7c). By Day 55, vehicle treated H1975-OR xenografts more than doubled in size to 473 ± 51 mm^3^, whereas NGI-189 treated tumors showed no significant progression from the beginning of treatment with an average volume of 190 ± 37 mm^3^ (p= 0.004). Remarkably, three NGI-189 treated tumors did not progress by 90 days, driving a significantly improved survival rate when compared to vehicle treated controls [60% vs. 0%; p=0.022 (Fig. 7d)]. Again, treatment with NGI-189 was well tolerated and mouse weight during treatment was comparable in vehicle or osimertinib treated mice (Fig. 7e).

For HCC827-GR xenografts, average tumor volumes for vehicle treated mice more than doubled to 411 ± 38 mm^3^ at Day 15. However, xenografts did not progress at this time point in the NGI-189 treated group with a significantly smaller average volume of 186 ± 16 mm^3^ (p<0.005, Fig. 7f). Although all HCC827-GR tumors ultimately progressed, survival rates were significantly longer with NGI-189 treatment (17.6 vs 23.7 days, p<0.005; Fig. 7g) and OST inhibitor therapy was well tolerated (Fig. 7h). Together, these preclinical tumor models demonstrate that partial inhibition of N-glycosylation with aminobenzamidosulfonamide small molecule OST inhibitors can be leveraged to induce tumor regression and to delay tumor growth of RTK driven NSCLC xenografts.

## DISCUSSION

In this work we demonstrate the feasibility for *in vivo* translation of a new therapeutic approach in oncology: partial inhibition of N-glycosylation. Systematic medicinal chemistry and analog screening efforts delivered small molecules with superior potency and were then leveraged to demonstrate inhibition of this PTM *in vivo* without significant toxicity. In EGFR mutant NSCLC, OST inhibition reduced EGFR dependent oncogenic signaling and proliferation *in vitro*, and caused tumor regression *in vivo*. Extensive characterization of aminobenzamidosulfonamide OST inhibitors also elucidated the mechanistic basis for partial inhibition of N-glycosylation, a biologic effect that has not previously been studied and underlies therapeutic efficacy in animal models of lung cancer.

N-glycosylation is an essential cellular process, and pharmacologic strategies to completely block this co- and post-translational protein modification are anticipated to effect both normal and transformed cells. Indeed, the toxicity of blocking all N-glycosylation is well exemplified by tunicamycin, a natural product that prevents synthesis of N-glycan precursors and is a potent mammalian toxin (*29, 38*). In contrast, aminobenzamidosulfonamide OST inhibitors produce partial inhibition of N-glycosylation. On a glycoproteomic level, NGI-1 is known to reduce transfer of N-glycans to proteins in a site-specific manner and with preferential skipping of NXS vs. NXT sequons (*39*). This finding, along with evidence for target engagement of the inhibitor with catalytically inactive or peptide binding site mutant STT3B, suggested that partial inhibition was caused by an allosteric mechanism of action that incompletely disrupts both OST-A and OST-B activity (*28, 32*). More recently, NGI-1 has been shown to act as an uncompetitive inhibitor that interacts with both the lipid linked glycan and the catalytic to disrupt the enzymatic reaction (*40*). Here we provide further mechanistic insight into how the inhibitors cause partial inhibition of N-glycosylation. By improving inhibitor potency by ∼10 fold and achieving complete inhibition of N-glycosylation in cells with only knockout of a single catalytic subunit, we establish that it is functional redundancy between the OST-A and OST-B complexes that prevents complete loss of N-glycosylation and confers tolerability.

OST targeting was conceived as an alternative and complementary approach for blocking RTK dependent signaling necessary for tumor growth in EGFR mutant NSCLC. The conspicuously high number of sequons encoded by the EGFR amino acid sequence (n=12) coupled with site directed mutagenesis studies showing loss of as few as four N-glycosylation sites impairs EGFR maturation and function (*41*), suggested this PTM as a vulnerability for disrupting receptor signaling. Because OST-A catalyzes glycosylation for the majority of sequons in the proteome, we hypothesized that optimization of inhibitors for this target would improve efficacy. However, we found that improved activity against either STT3A (with NGI-186, −189) or STT3B (with NGI-190) was effective in reducing EGFR stability and signaling. This insight has important implications for OST inhibitor drug development and underscores the value of deliberately optimizing inhibitory activity for an effector glycoprotein (i.e., EGFR) instead of a specific OST catalytic subunit. In this context, it is realistic to anticipate that that OST inhibitors can be further developed to maximize effects for additional N-glycosylated protein targets.

Inhibition of N-glycosylation, despite the partial effect due to the redundancy of OST-A and OST-B activities, will reduce glycan site occupancy for multiple ER translated proteins. To establish a causal relationship for inhibition of EGFR N-glycosylation, we developed an EGFR mutant NSCLC model addicted to an N-glycosylation independent EGFR chimera. This model system weighs the potential cellular effects of OST inhibitors on glycoproteins other than the EGFR in the setting of EGFR addiction. Our results show that CD8-EGFR-CL expression produces significant rescue of both lung cancer proliferation and clonogenic survival after treatment with OST inhibitors such as NGI-186, −189, and −190 and indicates the dominant effect of OST inhibition on cell viability in these tumor cells is caused by loss of EGFR function. In addition, although rescue was incomplete, the limited effects of OST inhibition on cell viability in the presence of CD8-EGFR-CL are consistent with the therapeutic window achieved *in vivo*.

From one perspective, the potential for effects on multiple downstream N-glycosylated target proteins could be considered a potential weakness for this therapeutic strategy. However, from another perspective, the ability to disrupt the function of multiple glycoproteins (and RTKs) can also be viewed as a distinctive strength. In EGFR mutant NSCLC, which frequently develops resistance to EGFR TKIs through EGFR kinase domain mutation or upregulation of parallel RTK signaling, OST inhibition can block the adaptive bypass signaling mediated by other glycoproteins. Herein we show that EGFR TKI resistance associated with upregulation of MET or PTK7 glycoproteins is also sensitive to OST inhibition, and given the emergence of TKI resistance and limited response to available immunotherapy-based interventions (*42, 43*), OST inhibition may provide a new therapeutic approach for this glycoprotein driven subset of NSCLC.

Establishing structure activity relationships for OST inhibitors enabled iterative pharmacologic optimization and identification of a new lead (NGI-189) which more potently prevents EGFR activation with corresponding reductions in downstream signaling through AKT and p70S6K. Previously we showed that OST inhibition with NGI-1 induced senescence (*27*), herein we demonstrate that the enhanced potency achieved with NGI-189 increases G1 cell cycle arrest with the appearance of sub-G1 populations and cell death markers such as Bim. Most notably, our medicinal chemistry efforts delivered a bioavailable inhibitor for *in vivo* testing and demonstrated that single agent NGI-189 treatment delayed or stopped tumor growth in multiple cell line derived or PDX tumors. In fact, all NGI-189 treated PDX or EGFR TKI resistant xenografts either regressed or did not progress at time points where controls at least doubled in size. The response of H1975-OR cells, which display slow growth kinetics *in vivo,* were of particular interest due to their remarkable sensitivity to OST inhibition and cure for 60% of treated animals. Whether OST inhibition has a more pronounced effects on tumors with a fitness deficit and slow growth remains to be determined and will require further investigation.

In summary we have leveraged medicinal chemistry efforts to define the pharmacology of OST inhibitors and to deliver potent and bioavailable small molecules like NGI-189 for development as therapeutic agents. Moreover, the demonstration of *in vivo* tolerability with partial inhibition of this PTM introduces unexplored territory for drug development. Notably, other mechanisms for partial loss of N-glycan site occupancy that affect N-glycosylation at sequons that differ from OST inhibition, such as loss of translocon associated protein (TRAP) complex function (*44*) or pharmacologic inhibition of the signal recognition particle receptor-*β* (*39*), have recently been discovered. Thus differential inhibition of OST catalytic subunit activity may be just the first pharmacologic approach for targeting N-glycosylation to alter protein function.

## SIGNIFICANCE

Enzymes that control oncogenic protein function through addition of post-translational modifications (PTM) are attractive targets for development of cancer therapeutics. N-glycosylation is a key PTM enzymatic reaction that is catalyzed in the lumen of the endoplasmic reticulum (ER) and is required for proper protein folding and trafficking. However, the pathway for synthesis and transfer of N-glycans is essential, with the potential for toxicity when blocked. Nevertheless, small molecule inhibitors of the oligosaccharyltransferase (OST), the ER membrane embedded enzyme complex that transfers glycans to newly synthesized proteins, have been demonstrated to partially reduce N-glycosylation; an observation that suggests tolerability and the potential for in vivo translation. Herein we advance the chemical biology of OST inhibitors and use them to dissect the mechanism underlying partial loss of N-glycosylation, a biological effect that was not previously known to be achievable. Through synthesis and testing of more than 200 inhibitor analogs that improved potency and other pharmacokinetic properties, we show that redundant activities of the mutually exclusive OST catalytic subunit paralogs (STT3A and STT3B) prevent the complete inhibition of N-glycosylation and confer tolerability. In addition, the results also demonstrate that OST inhibitor activities can be tailored to preferentially affect subsets of glycoproteins and provide a framework for OST drug development. In EGFR mutant NSCLC models in vitro, highly N-glycosylated RTKs are inactivated by OST inhibition whereas tumor cell survival is rescued by N-glycosylation independent EGFR signaling. Testing of a bioavailable inhibitor, NGI-189, was advanced to PDX and TKI resistant lung cancer models and demonstrated tumor regression without observable animal toxicity. Together this data suggests that cellular N-glycosylation can be pharmacologically manipulated to produce discrete cellular effects as well as anti-tumor activity in specific cancer subtypes.

## MATERIALS AND METHODS

### OST inhibitor synthesis

OST inhibitors were synthesized from readily available starting materials. The synthesis of NGI-186 is illustrated in Fig. S3 as a representative example and was completed in three steps. First, commercially available 5-(chlorosulfonyl)-2-fluorobenzoic acid (1) was reacted with morpholine and triethylamine in dichloromethane to give a sulfonamide intermediate (2). Subsequent coupling with 5-ethylthiazol-2-amine using HATU and diisopropylamine in DMF gave the corresponding amide (3). Finally, nucleophilic aromatic substitution of the fluoride with 1-cyclopropyl-N-methylmethylamine at 80°C afforded NGI-186 in 84% yield (4). A more detailed description of the synthetic methods used to prepare NGI-186, −189, and −190 are provided in the Suppl. Methods and Fig. S3.

Equilibrium solubility was measured in 0.2 M KH2PO4 aqueous buffer at pH 7.4 by Absorption Systems (Exton, PA). NGI-186 or −189 were dissolved in buffer and shaken overnight at room temperature. Samples were then centrifuged to remove insoluble material and the supernatant was diluted 1:1 in buffer:acetonitrile. Samples were then assayed by LC-MS/MS using electrospray ionization.

### Cell lines and pharmacological inhibitors

The H3255 (CVCL_6831), HCC-4006 (CVCL_1269), HCC-2935 (CVCL_1265), H1975 (CVCL_1511) and HCC827 (CVCL_2063) cells were purchased from ATCC. PC9 (CVCL_B260) cells were provided by Dr. Politi. The generation of Halo3N expressing HEK293 (CVCL_0063) WT, STT3A KO, and STT3B KO cells has been previously described (*28*). Derivation of the HCC827-GR and H1975-OR cells have also been described previously (*32, 37*). HEK293 cells were cultured in DMEM + 10% FBS (Gibco, Life Technologies, Grand Island, NY, US) + 0.5 µg/ml of G418 (Gibco, Life Technologies, Grand Island, NY, US). All NSCLC cells were cultured in RPMI 1640 + 10% FBS (Gibco, Life Technologies, Grand Island, NY, US), grown in a humidified incubator with 5% CO_2_, and kept in culture no more than 6 months after resuscitation from original stocks. Mycoplasma cell culture contamination was routinely checked and ruled out using the MycoAlert Mycoplasma Detection Kit (Lonza; Rockland, ME USA). Gefitinib and Osimertinib were purchased at Selleck Chemicals LLC (Houston, TX, USA). Paclitaxel was purchased at MedChem (Monmouth Junction, NJ USA). Luciferin was supplied by Promega (Madison, WI USA).

### Generation of CD8-EGFR-CL cell lines

Briefly, the CD8-EGFR was amplified as described (*35*) and two-point mutations (L858R and C797S) were introduced in the intracellular kinase domain using the QuickChange site-directed mutagenesis kit (Agilent, Santa Clara). Primer sequences are provided in Suppl. Methods and mutations were confirmed by DNA sequencing. The resulting plasmid with a CD8-EGFR transgene that harbors L858R and C797S mutations (CD8-EGFR-CL) was stably transfected into PC9 cells with Lipofectamine (Life Technologies) followed by selection with 0.7 µg/ml G418 and 1 µM osimertinib.

### Proliferation, clonogenic, and cell cycle analysis

Proliferation and growth rates were determined by MTT (Thiazolyl Blue Tetrazolium Bromide; Cat# M5655. Sigma-Aldrich, St. Louis, MO USA) according to the manufacturer’s directions. For each experiment, 1000 cells were seeded in triplicate wells in 96-well plates. The following day, cell cultures were treated with DMSO vehicle, EGFR TKI inhibitors, Paclitaxel, NGI-1, or OST inhibitor analogs as specified. Absorbance values were compared from day 5 to day 0. Clonogenic survival assays were performed under similar inhibitor experimental conditions and as described (*32*). Cell cycle distributions were analyzed following treatment with DMSO vehicle or 5 μM of NGI-1, NGI-186, or NGI-189 for 24 hours. Treated cells were trypsinized and centrifuged, washed once with ice-cold PBS, fixed with ice-cold 70% ethanol, and stored overnight at −20 °C. After washing twice with PBS, they were incubated for 30 minutes at room temperature in 200 μL of Guava Cell Cycle Reagent (Luminex, Boston, MA USA). Cytofluorometric acquisitions were performed on a LSRII cytometer (BD Biosciences). First line analysis was performed with FlowJo software 10.8.1 (Becton Dickinson & Company, Ashland, OR USA) upon gating of the events characterized by normal forward and side scatter parameters and discrimination of doublets in a PI-A vs. PI-W bivariate plot. Approximately 50,000 cells were analyzed per experiment.

### Western blot analysis

Primary antibodies are listed in Supplementary Table 3. The nitrocellulose-bound primary antibodies were detected with anti–rabbit IgG horseradish peroxidase–linked antibody or anti–mouse IgG horseradish peroxidase–linked antibody (EMD Millipore; Temecula, CA USA), and were detected by the enhanced chemiluminescence staining ECL (GE Healthcare–Amersham Pharmacia, Buckinghamshire, U.K.). Densitometry and EGFR shift of size were determined using Image J 1.53t (National Institutes of Health, USA) and normalized by each corresponding loading control.

### OST inhibitor imaging for bioavailability *in vivo*

Bioluminescent imaging of mice bearing MDA-MB-231 ER-LucT flank tumors was performed as previously described (*29*). Tumors were grown in 6-week-old female athymic Swiss nu/nu mice (Envigo, Indianapolis, IN USA) by subcutaneous flank implantation of approximately 1 × 10^7^ cells into the hind limb. Thirteen days following injection, base line bioluminescent imaging using an IVIS Spectrum (Perkin Elmer. Shelton, CT USA) was performed over a 30-minute time period and following a single *i.p.* dose of 150 mg/kg luciferin in normal saline using a temperature-controlled bed during image acquisition. Mice were then randomized to vehicle or 10mg/kg NGI-186 treatment groups and bioluminescence for regions of interest encompassing each tumor was measured at 6 and 24 hours to determine peak bioluminescent activity and calculate changes over time.

### OST inhibitor therapeutic studies in xenograft tumors

The effects of NGI-189 were evaluated in female immunodeficient NSG mice (YU-006 and YU-010 PDX. Jackson Laboratory) or athymic Swiss nu/nu mice (H1975-OR and HCC827-GR; Envigo) inoculated in flanks of 6-to-8-week-old mice. For PDX experiments, fresh-frozen tumor tissues were processed with a tumor dissociation kit (Cat# 130-095-929. Miltenyi Biotech, Waltham, MA USA) to generate a single cell suspension and then resuspended in Matrigel (Cat# 356237. Corning, NY USA). Tumor xenografts were established by injection of 1 × 10^6^ (PDX) or 5 × 10^6^ (cell lines) tumor cells subcutaneously into the right hind leg. When tumors reached ∼200 mm^3^ animals were randomized to receive *i.p.* NGI-189 (10 mg/kg) or vehicle for a total of 8 doses (See legends for treatment details). For H1975-OR tumors, osimertinib was orally administered every day at a dose of 5 mg/kg over the same time-period as NGI-189 treatment. Of note, one H1975-OR bearing mouse from the vehicle-treated group died unexpectedly at day 40 after cell injection and was removed from the survival analysis so that the magnitude of NGI-189 effects were not inflated by this event. Tumor size was measured 3 times per week, and volume was calculated according to the formula π/6 x (length) x (width)^2^. Tumor growth delay was evaluated based on the time required for tumor volume doubling. Data are expressed as the mean volume ± SEM tumor volume. Group sizes are specified in the respective figures.

For pathology studies, six nude mice were randomized to receive three doses of vehicle (n=3) or 10 mg/kg of NGI-189 (n=3) every other day. Twenty-four hours after the last dose, mice were sacrificed, and samples were collected. For immuno-fluorescence mutant EGFR with exon 19 deletion was evaluated using 5-um thick paraffin embedded tissue dewaxed and hydrated in xylene and ethanol, respectively. Antigen retrieval was performed in citrate buffer in a microwave at pH 6.0. The slides were permeabilized with 0.1% Triton X-100 PBS and blocked with 0.1% Triton X-100 PBS containing 10% goat serum and 10% horse serum (Cat# G9023 and Cat# H0146, respectively. Sigma-Aldrich, St. Louis, MO USA). The primary antibody was added at a 1:100 dilution overnight at 4°C followed by 1h incubation with Alexa Fluor 488-conjugated secondary antibody (1:500; Suppl. Table 1). The slides were then counter-stained with a mixture (1:250 dilution) of 4′,6-diamidino-2-phenylindole (DaPi. Cat# D9542. Sigma-Aldrich, St. Louis, MO USA) and bisbenzimide H 33342 trihydrochloride solution (Hoechst 33342. Cat# B2261. Sigma-Aldrich, St. Louis, MO USA), mounted, and visualized under the EVOS M5000 Fluorescent Microscope (AMF5000. Thermo Fisher Scientific Inc., Waltham, MA USA) with 20X objective. Ten random fields of viable tumor per slide were captured and evaluated for protein expression levels.

### Statistical analysis

Results are expressed as mean ± standard error (S.E.) unless otherwise indicated. The Microsoft Excel (version 16.51) was used for statistical data analysis. GraphPad v 10.1.0 (264) was used for the statistical data analysis to compare the survival curves (time to reach 500 mm^3^; Fig. 7) and for simple linear regression analysis (OST inhibition effect vs rescue; Fig. S1). Statistically significant differences in between-group comparisons were defined at a significance level of P-value ≤ 0.05 in 2-tail student’s t-Test of two-samples of unequal variance (heteroscedastic), Pearson’s R, or in Log-rank (Mantel-Cox) test.

### Study approval

All experimental procedures for animals were approved in accordance with Institutional Animal Care and Use Committee and Yale university institutional guidelines for animal care and ethics and guidelines for the welfare and use of animals in cancer research.

## ACKNOWLEDGEMENTS

This work was funded by the US National Institutes of Health (NIH) RO1CA240418 (J.N.C.), US National Institutes of Health (NIH) R41AI134531 (M.V), and by US National Institutes of Health (NIH) P50CA196530 and US National Institutes of Health (NIH) U01CA235747 (K.P.). We thank Yale Flow Cytometry for their assistance with and use of the LSR II flow cytometer. The Core is supported in part by an NCI Cancer Center Support Grant # NIH P30CA016359.

## AUTHOR CONTRIBUTIONS

Conception and design: M.B., C.J.Z., M.V., J.N.C.

Development of methodology: M.B., H.L., V.K., C.P., M.V., R.L., K.P., C.J.Z., J.N.C.

Acquisition of data: M.B., H.L., V.K., C.P., M.V., R.L., C.J.Z., J.N.C.

Analysis and interpretation of data: M.B., H.L., V.K., C.P., M.V., K.P., C.J.Z., J.N.C.

Writing original draft: M.B., J.N.C.

Writing, review, and/or revision of the manuscript: M.B., H.L., V.K., C.P., M.V., C.J.Z., K.P., J.N.C.

Study supervision: M.B., J.N.C.

## DATA AND MATERIAL AVAILABILITY

All data are available in the main text or the supplementary materials

