## Supplementary data and material and methods for "OST Catalytic Subunit Redundancy Enables Therapeutic Targeting of N-Glycosylation"

**The PDF file includes:**

Fig. S1 to S3
Table S1 to S3
Supplementary material and methods

**Other Supplementary materials for this manuscript includes the following:**

Unedited blots and gel images
Supporting data values file

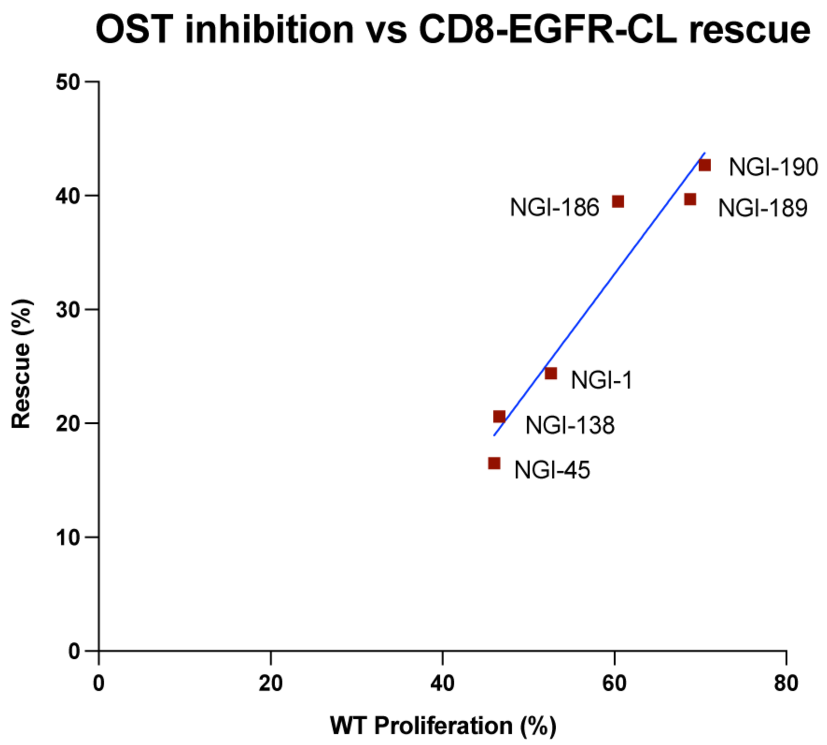


**Supplementary Figure 1. Correlation of OST inhibitor effect with CD8-EGFR-CL rescue.** Inhibition of proliferation (%) vs. rescue of proliferation (%) for six OST inhibitor analogs. Data points represent the average of three independent experiments. Simple linear regression analysis to calculate Pearson's R was used to assess statistical significance.

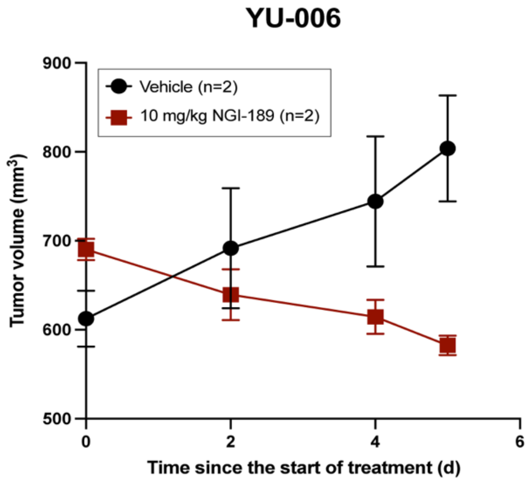


**Supplementary Figure 2. YU-006 tumors used for pathology.** YU-006 average xenograft tumor growth following treatment with vehicle or 10 mg/kg of NGI-189. Treatment was delivered by intraperitoneal injection of 3 doses every other day. Values are the mean ± SE of two tumors per group.

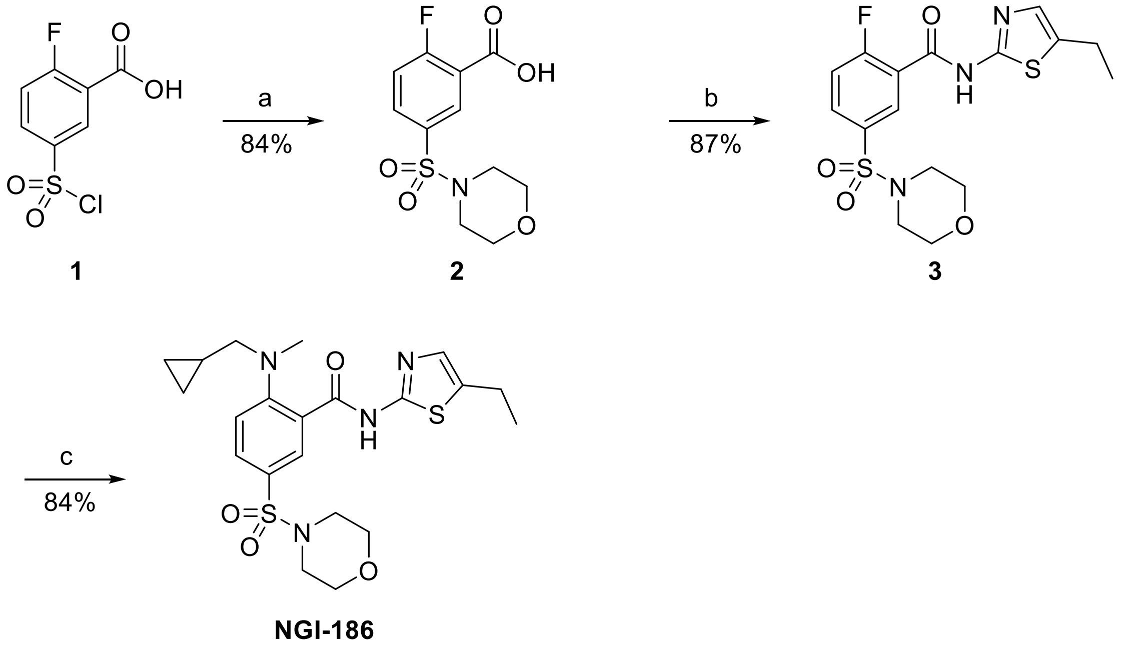


**Supplementary Figure 3. Reagents and reaction conditions for synthesis of NGI-186:** (a) morpholine, Et3N, CH2Cl2, 0-20° C, 40 h; (b) 5-ethylthiazol-2-amine, HATU, diisopropylethylamine, DMF, 12 h; (c) 1-cyclopropyl-N-methylmethylamne, 80°C, 6h. Percent yields for reaction conditons are reported.

**Supplementary Table 1. Mouse cytopathology analysis for NGI-189 analog.**

| Organ | Vehicle (n=3) | | | NGI-189 (n=3) | | |
|---|---|---|---|---|---|---|
| | 1 | 2 | 3 | 1 | 2 | 3 |
| Brain | NAD | NAD | NAD | NAD | NAD | NAD |
| Heart, lung, esophagus, trachea | NAD | NAD | 1* | NAD | NAD | NAD |
| Kidney, adrenal glands, liver, spleen, pancreas | NAD | 2* | 2* | NAD | NAD | 2*, 3* |
| Stomach, small and large intestine, mesenteric nodes | NAD | NAD | NAD | NAD | NAD | NAD |
| Skin, reproductive tract, urinary bladder, cervical lymphglands | NAD (female) | NAD (female) | NAD (female) | NAD (female) | NAD (female) | NAD (female) |
| Eye, Harderian gland | NAD | NAD | NAD | NAD | NAD | NAD |

NAD = No abnormality detected

1. Multifocal lung perivascular and peribronchial inflammation.
2. Rare renal cortex perivascular inflammation and glomerular sclerosis.
3. Rare small foci of perivascular liver inflammation.

*Minor abnormalities are consistent with age related changes.

**Supplementary Table 2. Mouse CBC and chemistry analysis for NGI-189 analog.**

| SuperChem | | | | | | | |
|---|---|---|---|---|---|---|---|
| Tests | Vehicle | | | NGI-189 | | | p-value |
| | mean | SEM | N | mean | SEM | N | |
| Total protein (g/dL) | 5.6 | 0.1 | 3 | 5.5 | 0.4 | 3 | 0.774 |
| Albumin (g/dL) | 3 | 0.1 | 3 | 3.1 | 0.1 | 3 | 0.496 |
| Globulin (g/dL) | 2.6 | 0.1 | 3 | 2.4 | 0.3 | 3 | 0.529 |
| A/G Ratio | 1.1 | 0.1 | 3 | 1.3 | 0.1 | 3 | 0.257 |
| AST (SGOT) (IU/L) | 70.3 | 18 | 3 | 112 | 55.1 | 3 | 0.536 |
| ALT (SGPT) (IU/L) | 24 | 3 | 3 | 31.7 | 7.3 | 3 | 0.409 |
| Alk Phosphatase (IU/L) | 70.7 | 9.5 | 3 | 72 | 6 | 3 | 0.913 |
| GGT (IU/L) | 1 | 0 | 3 | 1 | 0 | 3 | 1.000 |
| Total Bilirubin (mg/dL) | 0.2 | 0.1 | 3 | 0.1 | 0 | 3 | 0.686 |
| BUN (mg/dL) | 16.3 | 2.4 | 3 | 20 | 0.6 | 3 | 0.264 |
| Creatinine (mg/dL) | 0.2 | 0 | 3 | 0.2 | 0 | 3 | 1.000 |
| BUN/Creatinine Ratio | 81.7 | 12 | 3 | 100 | 2.9 | 3 | 0.264 |
| Phosphorus (mg/dL) | 8.4 | 0.4 | 3 | 9 | 0.7 | 3 | 0.529 |
| Glucose (mg/dL) | 249 | 15.5 | 3 | 239.7 | 19.9 | 3 | 0.731 |
| Calcium (mg/dL) | 10.6 | 0.2 | 3 | 10.4 | 0.2 | 3 | 0.577 |
| Magnesium (mEq/L) | 2.3 | 0.1 | 3 | 2.1 | 0.1 | 3 | 0.485 |
| Sodium (mEq/L) | 151 | 2.1 | 3 | 154.3 | 0.9 | 3 | 0.247 |
| Potassium (mEq/L) | 8.4 | 0.4 | 3 | 9.2 | 0.7 | 3 | 0.414 |
| NA/K Ratio | 18 | 0.6 | 3 | 17 | 1.5 | 3 | 0.590 |
| Chloride (mEq/L) | 110.3 | 0.7 | 3 | 113.3 | 1.9 | 3 | 0.243 |
| Cholesterol (mg/dL) | 107 | 3.2 | 3 | 102 | 9.6 | 3 | 0.664 |
| Triglyceride (mg/dL) | 116 | 16.5 | 3 | 101.7 | 10.7 | 3 | 0.512 |
| Amylase (IU/L) | 648.7 | 72.8 | 3 | 698.3 | 34.4 | 3 | 0.583 |
| PrecisionPSL (IU/L) | 39.7 | 10.2 | 3 | 40 | 5.9 | 3 | 0.979 |
| CPK (IU/L) | 192.3 | 36.2 | 3 | 436 | 362.8 | 3 | 0.572 |
| **Complete Blood Count** | | | | | | | |
| Tests | Vehicle | | | NGI-189 | | | p-value |
| | mean | SEM | N | mean | SEM | N | |
| WBC (10^3/uL) | 6 | 0.9 | 3 | 7.3 | 2.4 | 3 | 0.658 |
| RBC (10^6/uL) | 7.8 | 0.4 | 3 | 8.8 | 0.5 | 3 | 0.213 |
| HGB (g/dL) | 13 | 0.6 | 3 | 14.4 | 1.1 | 3 | 0.349 |
| HCT (%) | 43.7 | 2.4 | 3 | 48.7 | 2.3 | 3 | 0.210 |
| MCV (fL) | 56.3 | 2.8 | 3 | 55.7 | 2.7 | 3 | 0.874 |

| | | | | | | | |
|---|---|---|---|---|---|---|---|
| MCH (pg) | 16.9 | 0.2 | 3 | 16.3 | 0.5 | 3 | 0.388 |
| MCHC (g/dL) | 30 | 1.2 | 3 | 29.3 | 1.3 | 3 | 0.725 |
| Platelet Count (10^3/uL) | 714.7 | 260 | 3 | 940.3 | 207.3 | 3 | 0.536 |

**Differential (Absolute)**

| Tests | Vehicle | | | NGI-189 | | | p-value |
|---|---|---|---|---|---|---|---|
| | mean | SEM | N | mean | SEM | N | |
| Neutrophils (/uL) | 1298.7 | 118.7 | 3 | 1480.3 | 705.6 | 3 | 0.822 |
| Bands | NA | | 3 | NA | | 3 | |
| Lymphocytes (/uL) | 4231.3 | 713.3 | 3 | 4740.3 | 1563.2 | 3 | 0.788 |
| Monocytes (/uL) | 388.7 | 201.5 | 3 | 886.3 | 660.3 | 3 | 0.535 |
| Eosinophils (/uL) | 81.3 | 25.1 | 3 | 193 | 130.5 | 3 | 0.484 |
| Basophils (/uL) | 0 | 0 | 3 | 0 | 0 | 3 | 1.000 |

**Differential (Percentage)**

| Tests | Vehicle | | | NGI-189 | | | p-value |
|---|---|---|---|---|---|---|---|
| | mean | SEM | N | mean | SEM | N | |
| Neutrophils (%) | 22.7 | 3.5 | 3 | 22 | 6.8 | 3 | 0.936 |
| Bands (%) | 0 | 0 | 3 | 0 | 0 | 3 | 1.000 |
| Lymphocytes (%) | 70.3 | 1.5 | 3 | 66.7 | 4.5 | 3 | 0.506 |
| Monocytes (%) | 5.7 | 2.6 | 3 | 9 | 6.1 | 3 | 0.654 |
| Eosinophils (%) | 1.3 | 0.3 | 3 | 2.3 | 1.3 | 3 | 0.535 |
| Basophils (%) | 0 | 0 | 3 | 0 | 0 | 3 | 1.000 |

**Supplementary Table 3. List of antibodies used in the study.**

| Antibodies | dilution | source | identifier | application |
|---|---|---|---|---|
| **Rabbit monoclonal anti-HaloTag** | 1:3,000 | Promega (Madison, WI USA) | #G9211 | WB |
| **rabbit anti–EGFR** | 1:1,000 1:2,000 | Cell Signaling (Danvers, MA, USA) | #4267S | WB |
| **rabbit anti–pEGFR-Y1068** | 1:1,000 | Cell Signaling (Danvers, MA, USA) | #3777S | WB |
| **rabbit anti–AKT** | 1:1,000 | Cell Signaling (Danvers, MA, USA) | #9272S | WB |
| **rabbit anti–pAKT-S473** | 1:1,000 | Cell Signaling (Danvers, MA, USA) | #4060S | WB |
| **rabbit anti–P70 S6K** | 1:1,000 | Cell Signaling (Danvers, MA, USA) | #2708S | WB |
| **rabbit anti–phosphor P70 S6K T389** | 1:1,000 | Cell Signaling (Danvers, MA, USA) | #9205S | WB |
| **mouse anti–S6 ribosomal protein (RP)** | 1:1,000 | Cell Signaling (Danvers, MA, USA) | #2317S | WB |
| **rabbit anti–pS6RP S234-235** | 1:1,000 | Cell Signaling (Danvers, MA, USA) | #2211S | WB |
| **rabbit anti–BIM** | 1:1,000 | Cell Signaling (Danvers, MA, USA) | #2819S | WB |
| **rabbit anti–BIM** | 1:1,000 | Cell Signaling (Danvers, MA, USA) | #2933S | WB |
| **rabbit anti–PTK7** | 1:1,000 | Cell Signaling (Danvers, MA, USA) | #25618S | WB |
| **rabbit anti-STAT3** | 1:1,000 | Cell Signaling (Danvers, MA, USA) | #4904S | WB |
| **rabbit anti–pSTAT3 Y705** | 1:1,000 | Cell Signaling (Danvers, MA, USA) | #9145S | WB |
| **rabbit anti–MET** | 1:1,000 | Cell Signaling (Danvers, MA, USA) | #4560S | WB |
| **rabbit anti–pMET-Y1234/1235** | 1:500 | Cell Signaling (Danvers, MA, USA) | #3077S | WB |
| **rabbit phospho-IGF-I Receptor β (Tyr1135) (DA7A8)** | 1:1000 | Cell Signaling (Danvers, MA, USA) | #3918S | WB |
| **rabbit IGF-I Receptor β (D23H3)** | 1:1000 | Cell Signaling (Danvers, MA, USA) | #9750S | WB |
| **mouse anti-GADPH** | 1:10,000 | Proteintech (Rosemont, IL, USA) | 60004-1-Ig | WB |
| **Goat Anti-Rabbit IgG Antibody, F(ab')2, HRP conjugate** | 1:1,000 1:2,000 | EMD Millipore; Temecula, CA USA | AQ132P | WB |
| **Rabbit Anti-Mouse IgG Antibody, F(ab')2, HRP conjugate** | 1:1,000 1:2,000 | EMD Millipore; Temecula, CA USA | AQ160P | WB |
| **Rabbit anti-EGF Receptor (E746-A750del Specific)** | 1:100 | Cell Signaling (Danvers, MA, USA) | #2085S | IF |
| **Goat anti-Rabbit IgG (H+L) Highly Cross-Adsorbed Secondary Antibody, Alexa Fluor™ 488** | 1:500 | Invitrogen (Waltham, MA USA) | A-11034 | IF |

Keywords: WB, western blot; IF, immuno-fluorescence.

### SUPPLEMENTARY MATERIAL AND METHODS:

**Chemistry General Methods.** Proton nuclear magnetic resonance ($^1$H NMR) spectra were determined on Agilent DD2 400 MHz, 500 MHz or 600 MHz NMR spectrometers. Chemical shifts are reported in parts per million (ppm) downfield from tetramethylsilane (internal standard) with coupling constants in hertz (Hz). Multiplicity is indicated by the following abbreviations: singlet (s), doublet (d), triplet (t), quartet (q), multiplet (m), broad (br). Mass spectra were recorded using an Advion Expression CMS mass spectrometer coupled with an Agilent 1260 analytical HPLC. Intermediate products from all reactions (non-polar compounds) were purified by either flash column chromatography or medium pressure liquid chromatography using a Biotage Isolera One with SNAP cartridge unless otherwise indicated. Analytical purity for all final compounds was determined using an Agilent Eclipse XDB-Phenyl analytical HPLC column (3.5 $\mu$; 4.6 x 150 mm) with an ultraviolet detector. Thin-layer chromatography using glass-backed silica plates containing a fluorescent indicator (0.25 mm, Whatman, Merck) was used to monitor reactions. Chromatograms were visualized using ultraviolet illumination, exposure to iodine vapors, or by dipping in an aqueous potassium permanganate solution. All starting materials were used without further purification. All reactions were carried out under an atmosphere of dried nitrogen or argon. Compound names are derived from the structures using ChemDraw Ultra 14.0.

**NGI-186:** ***Preparation of 2-((cyclopropylmethyl)(methyl)amino)-N-(5-ethylthiazol-2-yl)-5-(morpholinosulfonyl)benzamide***

*Step 1: synthesis of 2-fluoro-5-(morpholinosulfonyl)benzoic acid (2)*

While under nitrogen, a solution of and 5-(chlorosulfonyl)-2-fluorobenzoic acid (2.39 g, 10 mmol) in dichloromethane (20 mL) was cooled to 0 °C and treated with triethylamine (2.78 mL, 20 mmol) and morpholine (0.88 mL, 10 mmol). After stirring for 2 days at room temperature, the reaction was concentrated, suspended in water and acidified with HCl (3N) until pH=4~5. Extraction with ethyl acetate afforded the title compound as a white solid acid (2.44 g, yield 84%).

*Step 2: synthesis of N-(5-ethylthiazol-2-yl)-2-fluoro-5-(morpholinosulfonyl)benzamide (3)*

While under nitrogen, a solution of 2-fluoro-5-(morpholine-4-sulfonyl)benzoic acid (1g, 3.46 mmol) and 5-ethyl-1,3-thiazol-2-amine (443 mg, 3.46 mmol) in DMF (5mL) was treated with diisopropylethylamine (0.57mL, 3.46 mmol) and HATU (1.32 g, 3.46 mmol). After stirring for 12 h at room temperature, the reaction was diluted with ethyl acetate, washed with water, dried over anhydrous magnesium sulfate, filtered, and concentrated. Purification by flash column (30-50% ethyl acetate in hexanes) gave the title compound as a white solid (1.2 g, yield 87%). $^{1}$H NMR (400 MHz, Chloroform-d) δ 8.53 (dd, J = 6.8, 2.4 Hz, 1H), 7.97 (ddd, J = 8.7, 4.6, 2.5 Hz, 1H), 7.41 (dd, J = 10.8, 8.7 Hz, 1H), 7.13 (t, J = 1.2 Hz, 1H), 3.73 (dd, J = 5.7, 3.7 Hz, 5H), 3.07 – 2.98 (m, 4H), 2.82 (qd, J = 7.5, 1.1 Hz, 2H), 1.32 (t, J = 7.5 Hz, 3H).

*Step 3: synthesis of 2-((cyclopropylmethyl)(methyl)amino)-N-(5-ethylthiazol-2-yl)-5-(morpholinosulfonyl)benzamide (NGI-186)*

A solution of *N*-(5-ethyl-1,3-thiazol-2-yl)-2-fluoro-5-(morpholine-4-sulfonyl)benzamide (360 mg, 0.901 mmol) and 1-cyclopropyl-*N*-methylmethanamine hydrochloride (109 mg,

0.901 mmol) in DMSO (2 mL) was treated with and diisopropylethylamine (0.30 mL, 1.80 mmol) and warmed to 80 °C with stirring for 6 h. After cooling to room temperature, the reaction was diluted with ethyl acetate, washed with water, dried over anhydrous magnesium sulfate, filtered, and concentrated. Purification by preparative thin layer chromatography using a mixture of dichloromethane, methanol and saturated aqueous ammonium hydroxide (20:1:0.1) gave the title compound as a white solid (350 mg, 84% yield). $^1$H NMR (400 MHz, Chloroform-d) δ 13.20 (s, 1H), 8.61 (d, J = 2.4 Hz, 1H), 7.87 (dd, J = 8.4, 2.4 Hz, 1H), 7.46 (d, J = 8.5 Hz, 1H), 7.14 (t, J = 1.2 Hz, 1H), 3.77 – 3.69 (m, 4H), 3.06 – 2.99 (m, 5H), 2.94 (d, J = 12.8 Hz, 6H), 2.81 (qd, J = 7.5, 1.2 Hz, 2H), 1.32 (t, J = 7.5 Hz, 3H), 1.09 – 0.84 (m, 1H), 0.58 – 0.46 (m, 2H), 0.11 (dt, J = 6.1, 4.8 Hz, 2H).

**NGI-189:** ***Preparation of 2-(cyclopentyl(methyl)amino)-N-(5-ethylthiazol-2-yl)-5-(morpholinosulfonyl)benzamide***

A solution of *N*-(5-ethyl-1,3-thiazol-2-yl)-2-fluoro-5-(morpholine-4-sulfonyl)benzamide (50 mg, 0.13 mmol) and N-methylcyclopentanamine hydrochloride (17 mg, 0.13 mmol) in DMSO (0.2 mL) was treated with and diisopropylethylamine (32 mg, 0.25 mmol) and warmed to 80 °C with stirring for 6 h. After cooling to room temperature, the reaction was diluted with ethyl acetate, washed with water, dried over anhydrous magnesium sulfate, filtered, and concentrated. Purification by preparative thin layer chromatography using a mixture of dichloromethane, methanol and saturated aqueous ammonium hydroxide (20:1:0.1) gave the title compound as a white solid (25 mg, 42% yield). $^1$H NMR (400 MHz, Chloroform-d) δ 8.71 (d, J = 2.3 Hz, 1H), 7.90 (dd, J = 8.4, 2.4 Hz, 1H), 7.54 (d, J =

8.5 Hz, 1H), 7.17 (s, 1H), 3.78 – 3.70 (m, 4H), 3.63 (s, 1H), 3.08 – 3.01 (m, 4H), 2.93 (s, 2H), 2.81 (qd, J = 7.5, 1.2 Hz, 2H), 2.10 – 1.40 (m, 6H), 1.42 – 1.27 (m, 3H).

**NGI-190:** ***Preparation of 2-(3,3-difluoropyrrolidin-1-yl)-N-(5-ethylthiazol-2-yl)-5-(morpholinosulfonyl)benzamide***

A solution of *N*-(5-ethyl-1,3-thiazol-2-yl)-2-fluoro-5-(morpholine-4-sulfonyl)benzamide (50 mg, 0.13 mmol) and 3,3-difluoropyrrolidine hydrochloride (18 mg, 0.13 mmol) in DMSO (0.2 mL) was treated with and diisopropylethylamine (32 mg, 0.26 mmol) and warmed to 80 °C with stirring for 6 h. After cooling to room temperature, the reaction was diluted with ethyl acetate, washed with water, dried over anhydrous magnesium sulfate, filtered, and concentrated. Purification by preparative thin layer chromatography using a mixture of dichloromethane, methanol, and saturated aqueous ammonium hydroxide (20:1:0.1) gave the title compound as a white solid (35 mg, 57% yield). $^{1}$H NMR (400 MHz, Chloroform-d) δ 7.93(d, J = 4.0 Hz, 1H), 7.75 (dd, J = 8.0, 4.0 Hz, 1H), 6.88 (d, J = 8.0 Hz, 1H), 6.54 (s, 1H), 3.73 – 3.70 (m, 4H), 3.67 – 3.61 (m, 4H), 3.05 – 2.97 (m, 4H), 2.75 (q, J = 6.7 Hz, 2H), 3.53 – 2.43 (m, 2H), 1.30 (t, J = 6.7 Hz, 3H).

**Primers for site directed mutagenesis of CD8-EGFR.**

1. L858R-F (5'→3'): GATCACAGATTTTGGGCGGGCCAAACTGCTGGG
2. L858R-R (5'→3'): CCCAGCAGTTTGGCCCGCCCAAAATCTGTGATC
3. C797S-F (5'→3'): CATGCCCTTCGGCAGCCTCCTGGACTATGTC
4. C797S-R (5'→3'): GACATAGTCCAGGAGGCTGCCGAAGGGCATG
